# The effect of absent blood flow on the zebrafish cerebral and trunk vasculature

**DOI:** 10.1101/2020.07.23.216192

**Authors:** Elisabeth Kugler, Ryan Snodgrass, George Bowley, Karen Plant, Jovana Serbanovic-Canic, Paul C. Evans, Timothy Chico, Paul Armitage

## Abstract

The role of blood flow is complex and context-dependent. In this study, we quantify the effect of the lack of blood flow on vascular development and compare its impact in two vascular beds, namely the cerebral and trunk vasculature, using zebrafish as preclinical model. We performed this by analysing vascular topology, endothelial cell number, apoptosis, and inflammatory response in animals with normal blood flow or absent blood flow. We find that absent blood flow reduced vascular area and endothelial cell number significantly in both examined vascular beds, but the effect is more severe in the cerebral vasculature. Similarly, while stereotypic vascular patterning in the trunk is maintained, intra-cerebral vessels show altered patterning. Absent blood flow lead to an increase in non-EC-specific apoptosis without increasing tissue inflammation, as quantified by cerebral immune cell numbers and nitric oxide. In conclusion, blood flow is essential for cellular survival in both the trunk and cerebral vasculature, but particularly intra-cerebral vessels are affected by the lack of blood flow, suggesting that responses to blood flow differ between these two vascular beds.

**Key points:**

- We here use zebrafish as a model to quantitatively assess the impact of the lack of blood flow in development and compare its impact in two vascular beds, namely the cerebral to trunk vasculature.
- In both vascular beds, vascular growth and endothelial cell number are reduced by lack of blood flow, with increasing effect size from 2-5 days post fertilisation.
- Examination of vascular patterning shows that while stereotypic patterning in the trunk is preserved, the intra-cerebral vasculature patterning is altered.
- We found non-EC-specific cell death to be increased in both vascular beds, with a larger effect size in the brain, but that this cell death occurs without triggering tissue inflammation.

## Introduction

Endothelial cells (ECs) perform multiple functions during normal physiology including wound healing, tissue regeneration, immune response, menstruation, and pregnancy (Demir et al., 2010; Jung and Kleinheinz, 2013; Singer and Clark, 1999). Cerebral EC dysfunction is associated with neurodegenerative diseases, arteriovenous malformations, aneurysms, and stroke (Feigin Valery L. et al., 2017; Lackland and Weber, 2015). Increasing evidence suggests ECs display different molecular and functional properties according to their anatomical site such as the cerebral or trunk vessels (Abbott et al., 2010; Huntley et al., 2014; Kugler et al., 2019b; Vanlandewijck et al., 2018). This suggests novel insights could be obtained by studying the differential responses of these vascular beds to experimental manipulations.

Zebrafish are a frequently used model organism to study vascular development and disease (Bakkers, 2011; Chico et al., 2008; Gut et al., 2017). Fluorescent transgenic reporter lines allow cellular and sub-cellular visualization *in vivo* (Lawson and Weinstein, 2002). Advanced microscopy such as light sheet fluorescence microscopy (LSFM) acquires vascular information in greater anatomical depth and over prolonged periods of time (Huisken et al., 2004), allowing data acquisition to be rich in information and detail. Zebrafish embryonic transparency allows non-invasive and *in vivo* studies of different vascular beds in the same animal.

After 24 hours post fertilization (hpf) the zebrafish basic body plan is established and cardiac contraction starts. This is accompanied by vasculogenesis of primary vessels (Isogai et al., 2001, 2003). Blood flow plays an important role in processes such as EC polarization, vascular lumenisation, pruning, and heart trabeculation (Campinho et al., 2020; Chen et al., 2012; Hove et al., 2003; Kochhan et al., 2013; Lee et al., 2016; Lenard et al., 2015; Ricard and Simons, 2015), as well as directly impacts vascular architecture (Hoefer et al., 2013). Zebrafish embryos can survive for 7 days post fertilisation (dpf) without blood flow via oxygen diffusion due to their small size (Gut et al., 2017; Stainier and Fishman, 1994). This makes them well-suited to examine the role of blood flow on vasculogenesis and angiogenesis.

One approach to study the effect of absent blood flow in zebrafish is use of a cardiac troponin T2A (*tnnt2a)* morpholino (MO), which inhibits development of cardiac contraction, and thus blood flow (Sehnert and Stainier, 2002; Sehnert et al., 2002). The *tnnt2a* MO phenocopies the silent heart (*sih*) mutation (Sehnert et al., 2002). Additionally, control morpholino group account for injection-induced developmental delays (Stainier et al., 2017). Previous studies used this method to study the impact of blood flow on specific cerebral (Fortuna et al., 2015; Rödel Claudia Jasmin et al., 2019) and trunk vessels (Watson et al., 2013), showing reduced trunk EC numbers when blood flow is absent (Serbanovic-Canic et al., 2017; Watson et al., 2013).

Even though these studies have provided invaluable insights into the role of blood flow, important questions remain unanswered about the role of blood flow in vascular development. These include whether absent blood flow induces the same effects in different vascular beds and the effect of absent blood flow over time.

To examine these questions we here use LSFM 3D *in vivo* data acquisition in the head and trunk vasculature of 2-5dpf zebrafish embryos with and without blood flow. Our data show that even though the gross vascular responses to blood flow are comparable in different territories, some differences between anatomical sites exist, suggesting vascular territories exhibit differential sensitivity to absent blood flow.

## Results

### Cerebrovascular patterning is impaired by absent blood flow

We first examined whether cerebrovascular patterning is altered by lack of blood flow. Thus, we compared the cerebral vascular morphology of uninjected controls (**Fig. 1A-D**), control MO (**Fig. 1E-H**), and *tnnt2a* MO-injected embryos (**Fig. 1I-L**). The *tnnt2a* morphants were smaller with overtly abnormal cerebral vasculature, with effects becoming more severe from 2-to-5dpf. Intra-cerebral vessels, especially in the midbrain, were most severely affected showing altered growth patterns and an overall reduction of cerebral size was observed (**Fig. 1L**, dotted line). The primary head sinus (PHS), which extends laterally to the otic vesicle (OV), was enlarged, suggesting OV enlargement (**Fig. 1L**, white arrowheads), while peripheral/perineural vessels such as the primordial hindbrain channel (PHBC) were formed relatively normally during the investigated time frame (**Fig. 1L**, unfilled arrowhead).

**Figure 1.**
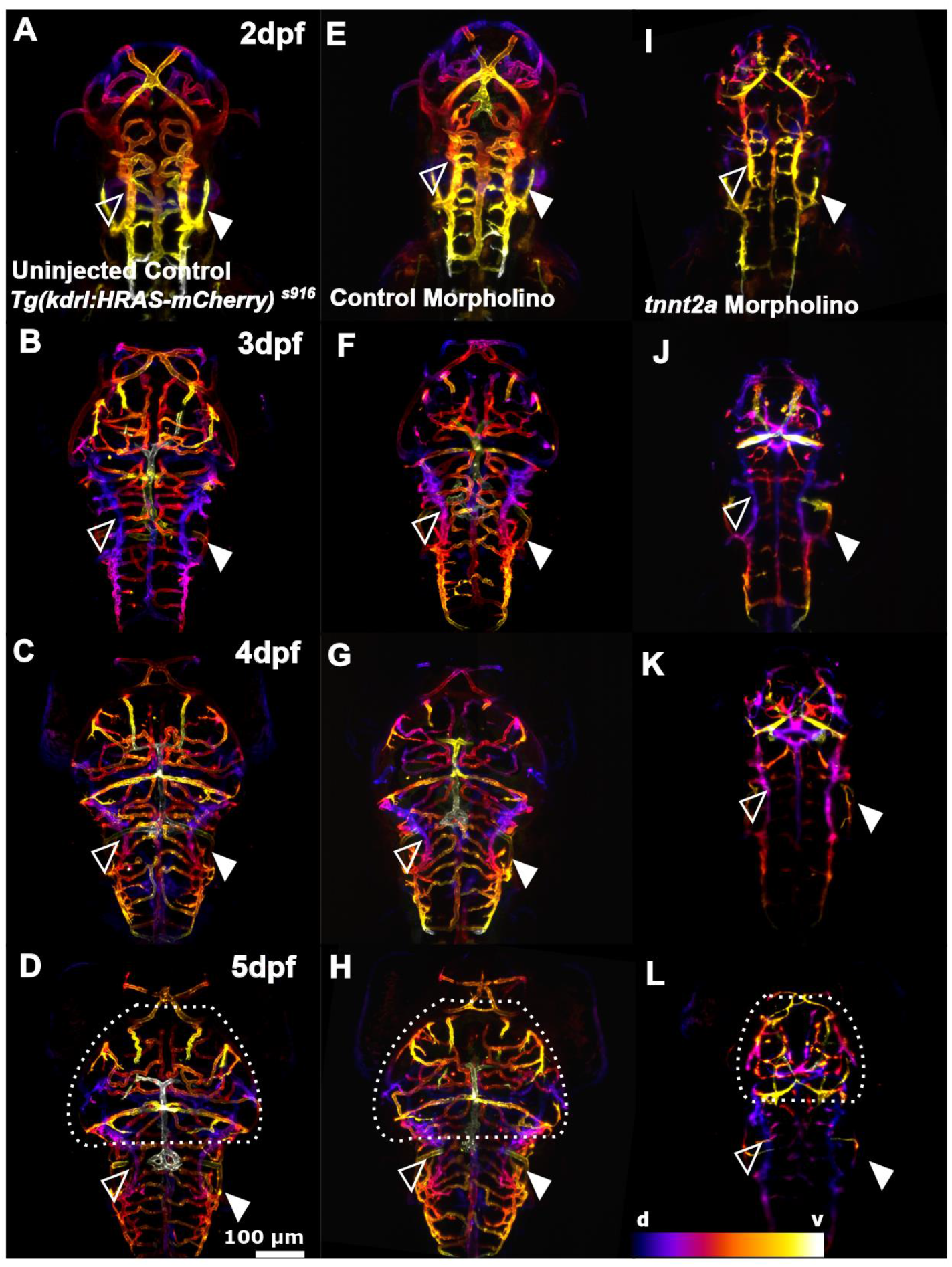
The effect of *tnnt2a* morpholino knockdown on cerebral vessel development. **(A-D)** Cerebral vasculature from 2-to-5dpf in *Tg(kdrl:HRAS-mCherry)*^*s916*^ uninjected control. **(E-H)** Cerebral vasculature in control MO. **(I-L)** Cerebral vasculature in *tnnt2a* MO injected samples (n=7-10; 2 experimental repeats). The lack of blood flow leads to a more severe vascular phenotype over time. Comparison between treatment groups shows that the midbrain vasculature is severely affected in *tnnt2a* MO (dotted lines) and the PCV surrounding the OV is enlarged (white arrowhead), while the PHBC appears normal (unfilled arrowhead; d – dorsal, v – ventral; representative images colour-coded by depth).

We next examined the distribution of EC in these animals (**Fig. 2**). This also suggested that peripheral vessels were less severely affected by absent blood flow, for example when comparing nuclei distribution in the middle cerebral vein (MCeV, **Fig. 2**, unfilled arrowhead) to central arteries (**Fig. 2**, white arrowhead). This was confirmed when examining Voronoi diagrams of EC nuclei distribution (**Fig. 2M-O**), which aid the visualization of EC nuclei density by partitioning MIPs into sub-regions based on nuclei position. Thus, visualising the spatial relationship of nuclei. This showed EC density to be unaltered in *tnnt2a* MO (white arrowhead), but reduced in the basilar artery (BA) or midbrain vessels.

**Figure 2.**
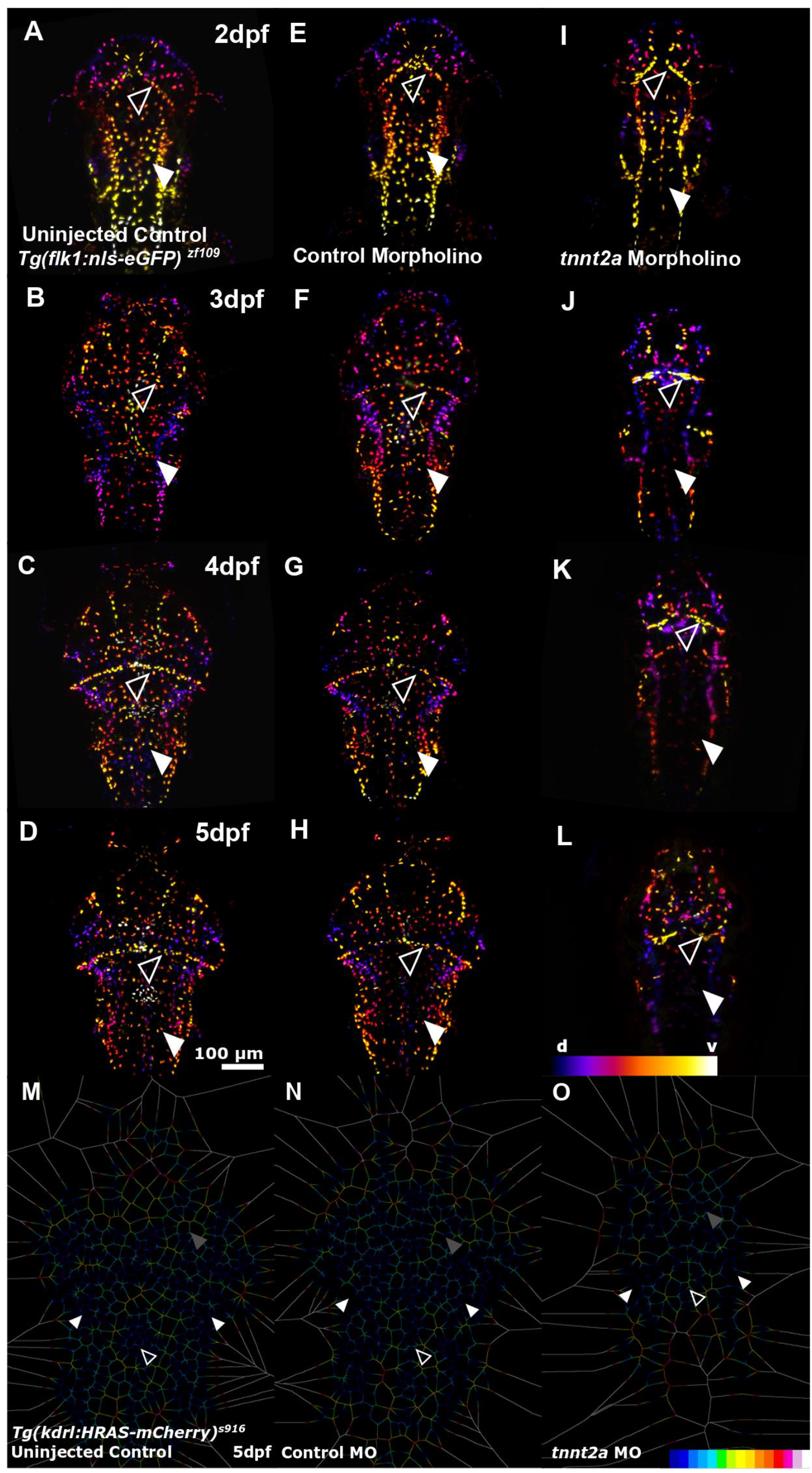
Lack of blood flow impacts cerebral EC number. **(A-D)** Cerebral EC nuclei from 2-to-5dpf in *Tg(flk1:nls-eGFP)*^*zf109*^ uninjected control. **(E-H)** Cerebral EC nuclei in control MO. **(I-L)** Cerebral EC nuclei in *tnnt2a* MO (n=7-10; 2 experimental repeats). Comparison of cerebral EC nuclei shows reduced cell numbers in *tnnt2a* MO with CtAs (white arrowhead) being particularly affected (d – dorsal, v – ventral; representative images colour-coded by depth). **(M-O)** Voronoi (image partitioning based on nuclei position) diagrams of cerebral EC nuclei suggests nuclei numbers to be maintained in peripheral vessels such as the PHBC (white arrowheads), while EC density is reduced in the midbrain (grey arrowhead) and BA (unfilled arrowhead).

This suggested that the effect of lacking blood flow becomes more severe over time and that central vessels are more impacted than peripheral vessels.

### Vascular patterning in the trunk is preserved, but vascular morphology altered

We next examined vascular patterning in the trunk of the same animals to study whether the observed effects were conserved or different in these vascular beds (**3A-L**). In contrast to the cerebral vessels, trunk vessel patterning was largely unaltered in the absence of flow. However, the morphology of the cardinal vein (CV) was less defined upon lack of blood flow (**Fig. 3**, white arrowhead).

**Figure 3.**
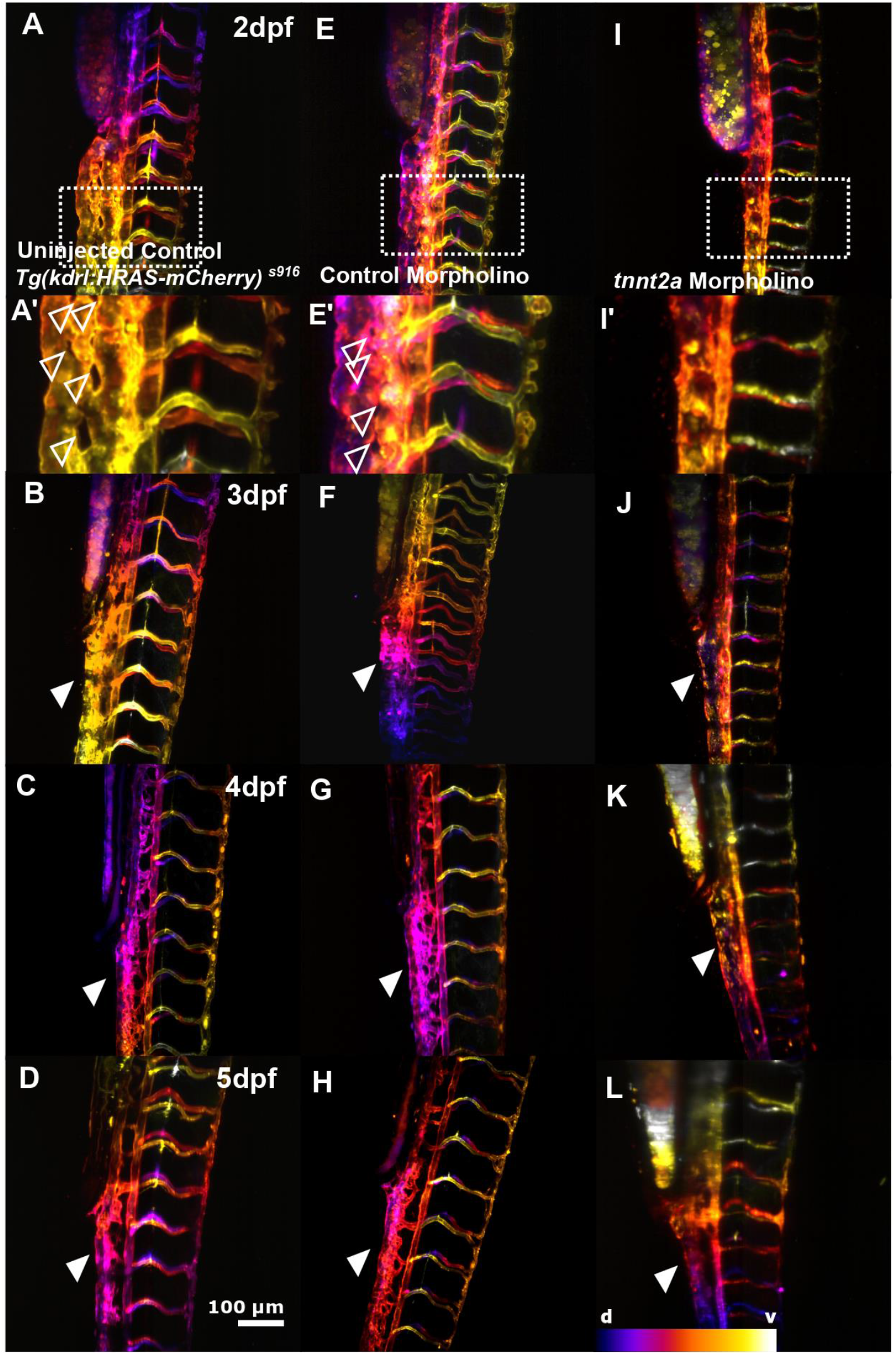
The effect of absent blood flow on trunk vessel development. **(A-D)** Trunk vasculature from 2-to-5dpf in *Tg(kdrl:HRAS-mCherry)*^*s916*^ uninjected control. **(E-H)** Trunk vasculature in control MO (n=7-10; 2 experimental repeats). **(I-L)** Trunk vasculature in *tnnt2a* MO. Topology of the trunk vasculature is established in the absence of flow, but morphology of the CV is severely altered and intussusceptions are lacking (unfilled arrowheads; arrowhead; d – dorsal, v – ventral; representative images colour-coded by depth).

Examination of EC nuclei suggested EC numbers in the trunk were less severely impacted by the lack of blood flow in comparison to the cerebral vessels (**Fig. 4**), with EC distribution in animals without blood flow being similar in the CV but altered in intersegmental vessels (ISVs) (**Fig. 4M-O**).

**Figure 4.**
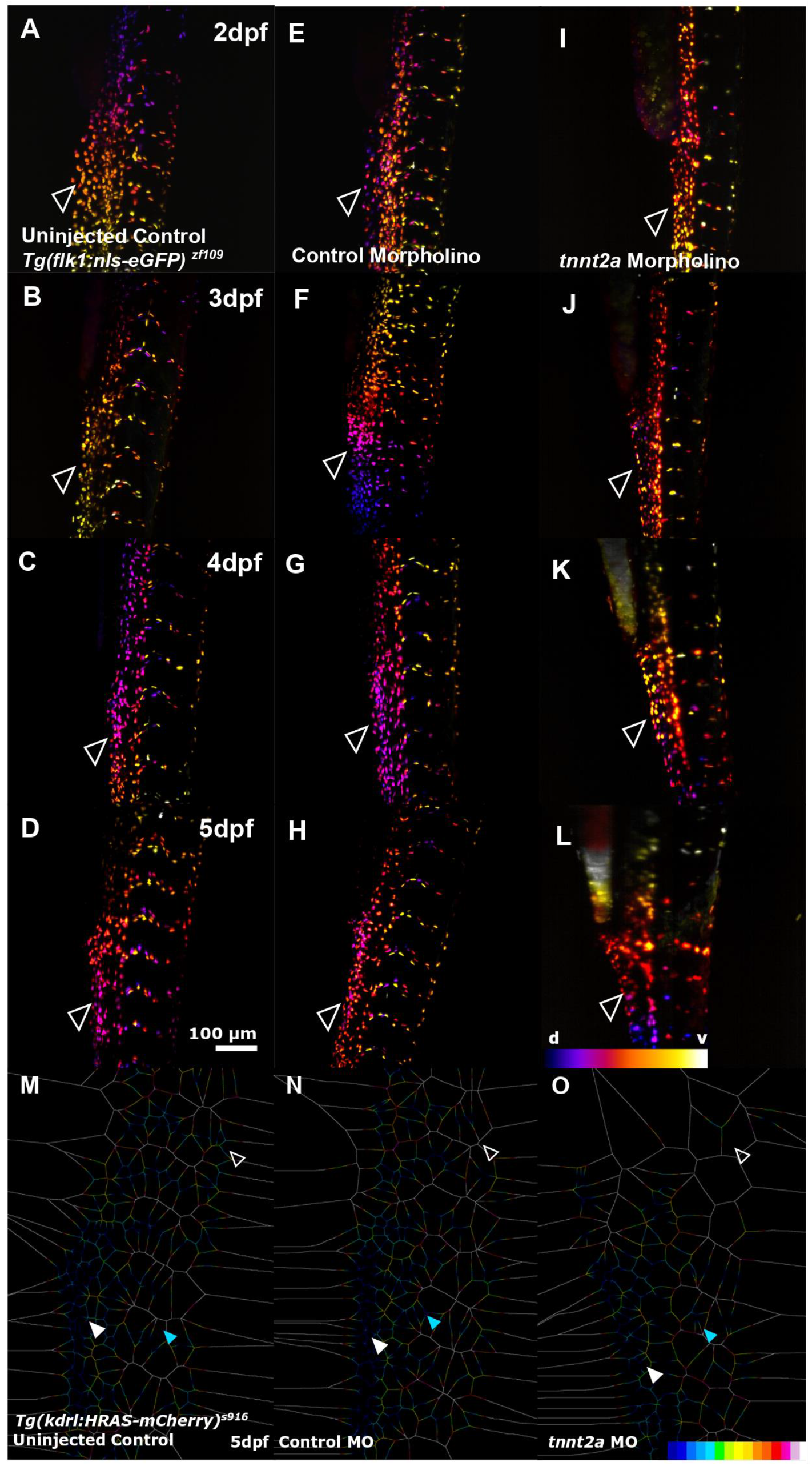
Lack of blood flow impacts trunk EC number. **(A-D)** Trunk EC nuclei from 2-to-5dpf in *Tg(flk1:nls-eGFP)*^*zf109*^ uninjected control. **(E-H)** Trunk EC nuclei in control MO. **(I-L)** Trunk EC nuclei in *tnnt2a* MO (n=7-10; 2 experimental repeats). Visual comparison of trunk EC nuclei suggests comparable numbers between treatment groups (d – dorsal, v – ventral; representative images colour-coded by depth). **(M-O)** Voronoi analysis of trunk EC nuclei shows EC distribution is maintained in the CV (white arrowhead), a decrease is observed in ISVs (blue arrowhead), and significant changes are observed more anteriorly (unfilled arrowhead).

Together, this suggested that vascular growth patterns in the trunk were unaltered, but remodelling, including intussusception (Karthik et al., 2018), were lacking.

### Lack of blood flow affects vascular area and number

To elucidate the effect of absent blood flow over time, we quantified cerebrovascular area, finding a significant reduction in *tnnt2a* morphants from 2-to-5dpf (**Fig. 5A**; uninjected control 2dpf p<0.0001, 3dpf p<0.0001, 4dpf p<0.0001, 5dpf p<0.0001). Similarly, the number of cerebral EC nuclei was significantly reduced in *tnnt2a* morphants from 2-to-5dpf (**Fig. 5B**; uninjected control 2dpf p 0.0409, 3dpf p<0.0001, 4dpf p<0.0001, 5dpf p<0.0001). To examine the relationship between vascular area and EC nuclei number, vascular area-to-nuclei ratios were calculated (**Fig. 5E**), showing a decrease in uninjected controls (0.78 fold-change 2-to-5dpf) and control MO (0.87 fold-change 2-to-5dpf), while the ratio increased in *tnnt2a* MO (1.69 fold-change 2-to-5dpf). This showed that the number of nuclei-to-vasculature increased in controls, but decreased in animals without blood flow.

**Figure 5.**
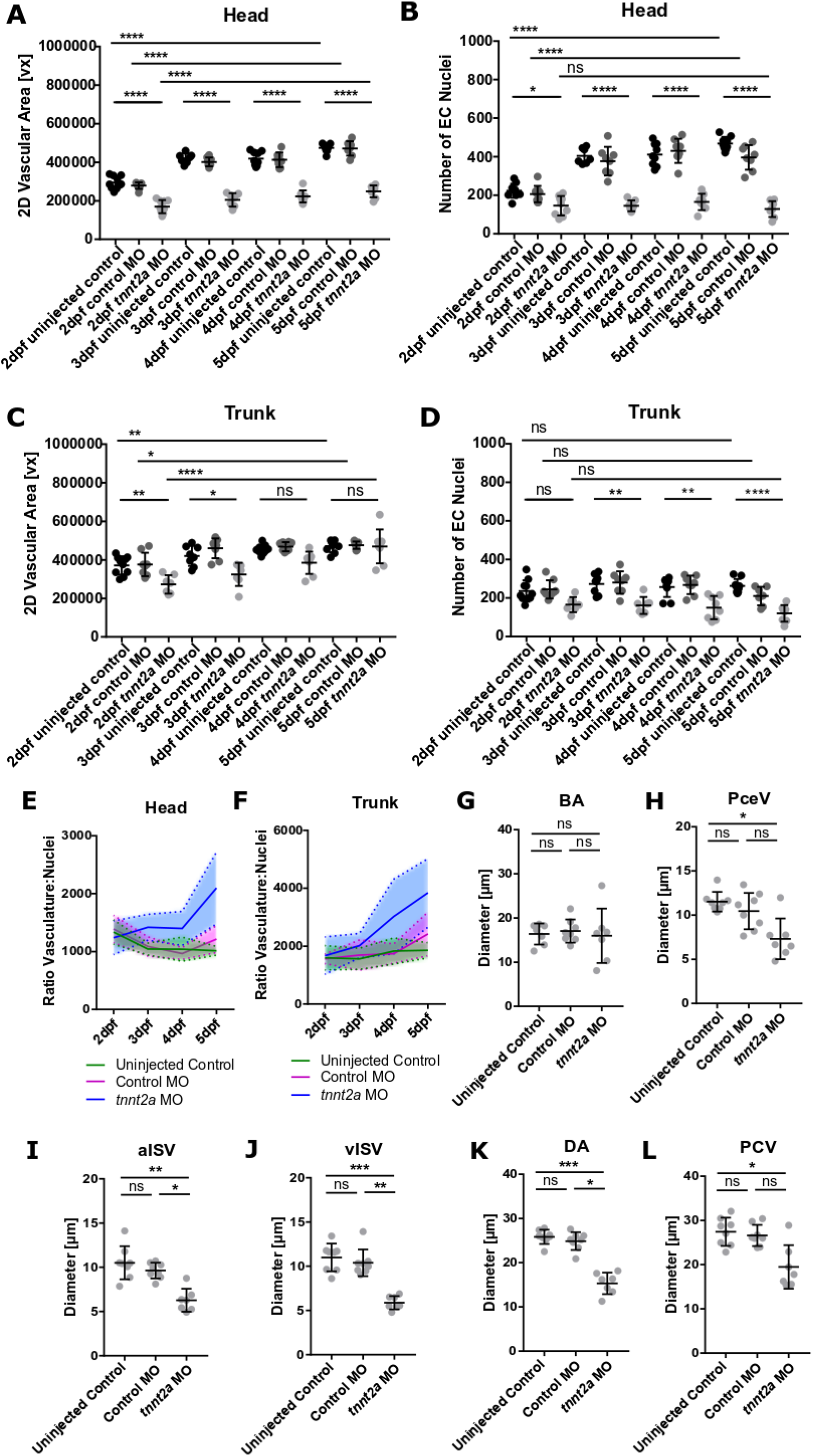
Quantification of vasculature and nuclei. **(A)** The cerebral vasculature is reduced in *tnnt2a* MO from 2-5dpf (n=7-10; 2 experimental repeats). **(B)** The number of cerebral EC nuclei is reduced in *tnnt2a* MO from 2-5dpf. **(C)** The trunk vasculature is reduced in *tnnt2a* MO at 2-3dpf, but not 4-5dpf (n=7-10; 2 experimental repeats). **(D)** The number of trunk EC nuclei is not altered in *tnnt2a* MO at 2dpf, but significantly reduced from 3-5dpf. **(E)** The ratio of cerebral vasculature-to-nuclei remains consistent in uninjected controls (green) and control MO (magenta), but increases over time in *tnnt2a* MO (blue). **(F)** The ratio of trunk vasculature-to-nuclei remains consistent in uninjected controls (green) and control MO (magenta), but increases over time in *tnnt2a* MO (blue). **(G)** The BA diameter was not changed upon lack of blood flow in comparison to uninjected controls (p>0.9999) and control MO (p>0.9999; Kruskal-Wallis test; n=7-9; 2 experimental repeats; 3dpf). **(H)** The PCeV diameter was reduced in comparison to uninjected controls (p 0.0175) but not control MO (p 0.1230; Kruskal-Wallis test). **(I)** aISV diameter was reduced in comparison to uninjected controls (p 0.0017) and control MO (p 0.0121; Kruskal-Wallis test). **(J)** vISV diameter was reduced in comparison to uninjected controls (p 0.0010) and control MO (p 0.0094; Kruskal-Wallis test). **(K)** The DA diameter was reduced in comparison to uninjected controls (p 0.0008) and control MO (p 0.0105; Kruskal-Wallis test). **(L)** The PCV diameter was reduced in comparison to uninjected controls (p 0.0159) but not control MO (p 0.0548; Kruskal-Wallis test).

We performed the same analysis on the trunk, and found that vascular area was significantly reduced by absent blood flow at 2dpf (p 0.0074) and 3dpf (p 0.0195), but not 4dpf (p 0.1904) and 5dpf (p>0.9999) in *tnnt2a* MO (**Fig. 5C**). Additionally, the number of nuclei was not significantly lower at 2dpf (p 0.1514; **Fig. 5D**) but was significantly decreased at 3-to-5dpf (3dpf p 0.0018, 4dpf p 0.0013, 5dpf p<0.0001). Analysis of vascular-to-nuclei ratio showed that vascular area increased in all three groups (**Fig. 5F;** fold-change 2-to-5dpf: uninjected control 1.10, control MO 1.55, *tnnt2a* MO 2.51).

Together, this suggested that the cerebral vessels were more severely affected than trunk vessels by absence of blood flow and that lack of blood flow increases vascular-to-nuclei ratio in both, the brain and trunk.

To study whether vessels of different identity were differentially or similarly impacted by the lack of blood, we quantified the diameter of selected arteries and veins in the brain and trunk 3dpf. Quantification of the cerebral BA diameter showed no difference between *tnnt2a* MO and controls (**Fig. 5G**; uninjected control p>0.9999, control MO p>0.9999), while the diameter of the posterior cerebral vein (PCeV) was reduced in animals without blood flow (**Fig. 5H**; uninjected control p 0.0175, control MO p 0.1230). Comparison of mean values showed that the BA diameter was reduced by 2.5% (uninjected control 16.41μm; *tnnt2a* MO 16μm), while the PCeV diameter was reduced by 36.46% (uninjected control 11.52μm; *tnnt2a* MO 7.32μm). Quantifying trunk ISV diameters, aISV diameter (**Fig. 5I**; uninjected control p 0.0017, control MO p 0.0121) and vISV diameter (**Fig. 5J**; uninjected control p 0.0010, control MO p 0.0094) were both reduced in *tnnt2a* MO. The mean diameter of aISVs was reduced by 40.4% (uninjected control 10.52μm; *tnnt2a* MO 6.27μm), while the mean diameter of vISVs was reduced by 46.64% (uninjected control 11μm; *tnnt2a* MO 5.87μm). The diameter of the dorsal aorta (DA) (**Fig. 5K**; uninjected control p 0.0008, control MO p 0.0105) as well as the posterior cardinal vein (PCV) (**Fig. 5L**; uninjected control p 0.0159, control MO p 0.0548) were reduced in *tnnt2a* MO. The mean diameter of the DA was reduced by 40.76% (uninjected control 25.86μm; *tnnt2a* MO 15.32μm), while the mean diameter of the PCV was reduced by 29.1% (uninjected control 27.46μm; *tnnt2a* MO 19.47μm).

Together, this suggests that vessels of arterial and venous identity are both affected by the lack of blood flow, but that vascular bed or vessel-specific differences might exist, as exemplified by the unchanged BA diameter.

### Non-EC-specific cell death is increased by absent blood flow

To examine whether cell death contributed to the reduced EC numbers observed in *tnnt2a* morphants, we quantified cell death using the live dye Acridine Orange.

Visual inspection of Acridine Orange levels suggested an overall increase in cell death in *tnnt2a* MO compared to controls in both the head (**Fig. 6A-C**) and trunk (**Fig. 6D-F**). However, specific foci of Acridine Orange were found in the brain of *tnnt2a* MO (white arrowhead). To examine whether the observed increase in Acridine Orange level were vascular or non-vascular, we extracted non-vascular (**Fig. 6A’-F’**) from vascular (**Fig. 6A’’-F’’**) signal by producing vascular masks in 3D, finding that the observed foci were non-vascular.

**Figure 6.**
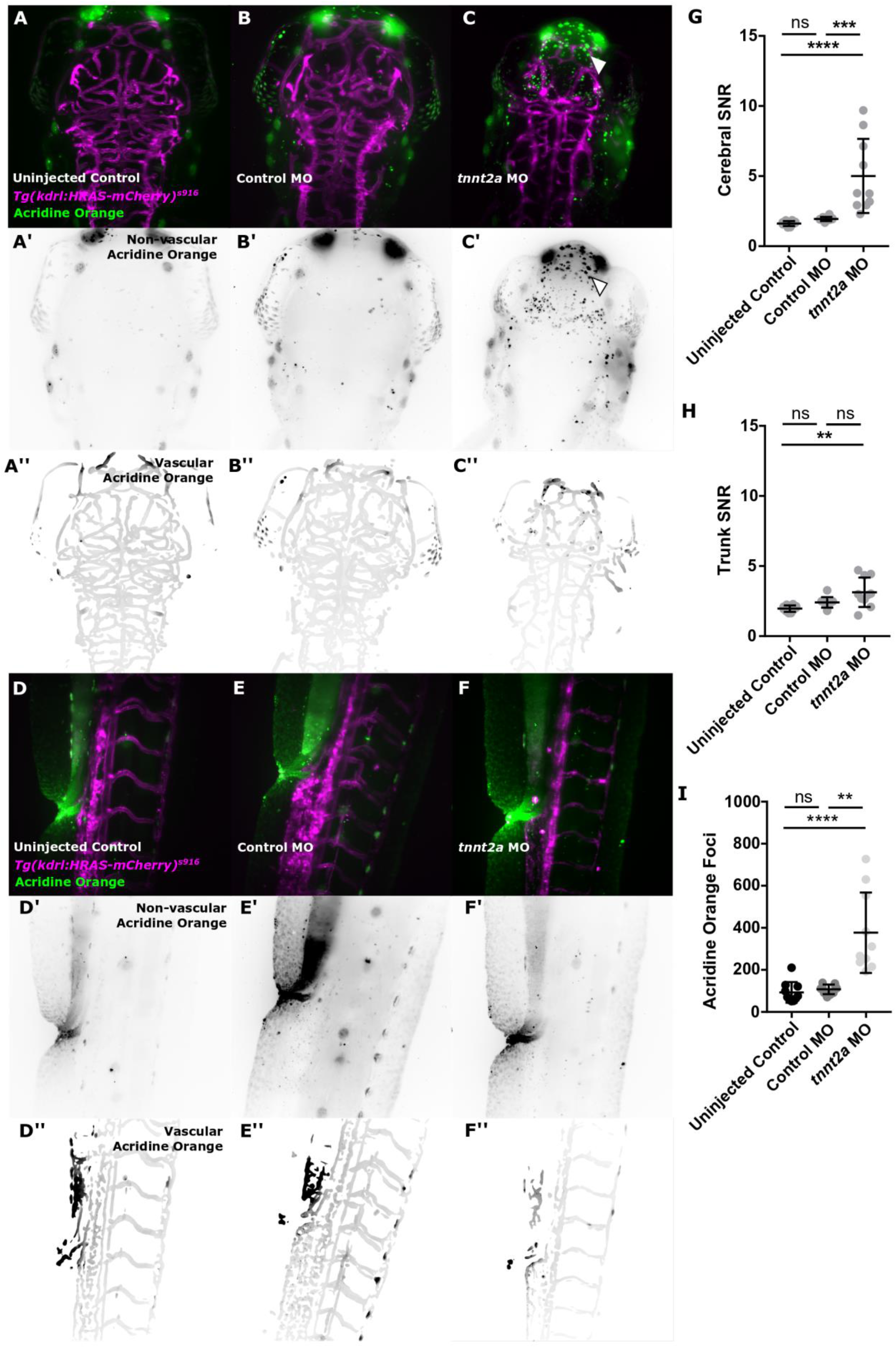
Cell death is increased by absent blood flow. **(A-C)** Cell death, visualized with Acridine Orange, is increased in *tnnt2a* MO in the cerebral vasculature (uninjected control n=10, control MO n=10, *tnnt2a* MO n=10; 2 experimental repeats; 3dpf). **(D-F)** Acridine Orange levels in the trunk vasculature appeared visually similar between groups. **(G)** Quantification of cerebral Acridine Orange SNR showed an increase in *tnnt2a* MO in comparison to uninjected controls (p<0.0001) and control MO (p 0.0003; One-Way ANOVA). **(H)** Quantification of trunk Acridine Orange SNR showed an increase in *tnnt2a* MO in comparison to uninjected controls (p 0.0014) but not control MO (p 0.0517; One-Way ANOVA). **(I)** Quantification of cerebral Acridine Orange foci showed an increase in *tnnt2a* MO in comparison to uninjected controls (p<0.0001) and control MO (p 0.0030; Kruskal-Wallis test).

Quantification of signal-to-Noise ratio (SNR; **Fig. 6G,H**) showed an increase in the head in *tnnt2a* MO compared with uninjected controls (p<0.0001, **Fig. 6G**) and trunk vasculature (p 0.0014, **Fig. 6H**). Additionally, the number of foci was increased in *tnnt2a* MO (**Fig. 6I**; uninjected control p<0.0001).

Together, this suggests that cell death is increased in *tnnt2a* MO, but that this is non-EC-specific.

### Inflammatory responses are not triggered by the absence of blood flow

We next examined whether immune cell numbers would be altered due to the observed cell death triggering an inflammatory response. As the observed effects of lack of blood flow were more severe in the head vasculature, quantification was only performed in this vascular bed.

Quantification of the number of macrophages in controls and *tnnt2a* MO at 3dpf (**Fig. 7A-C**) showed no difference (**Fig. 7D**; p 0.2356). Similarly, no difference was found in intracranial neutrophil numbers at 3dpf when comparing *tnnt2a* MO to controls (**Fig. 7E-G**; p 0.1708).

**Figure 7.**
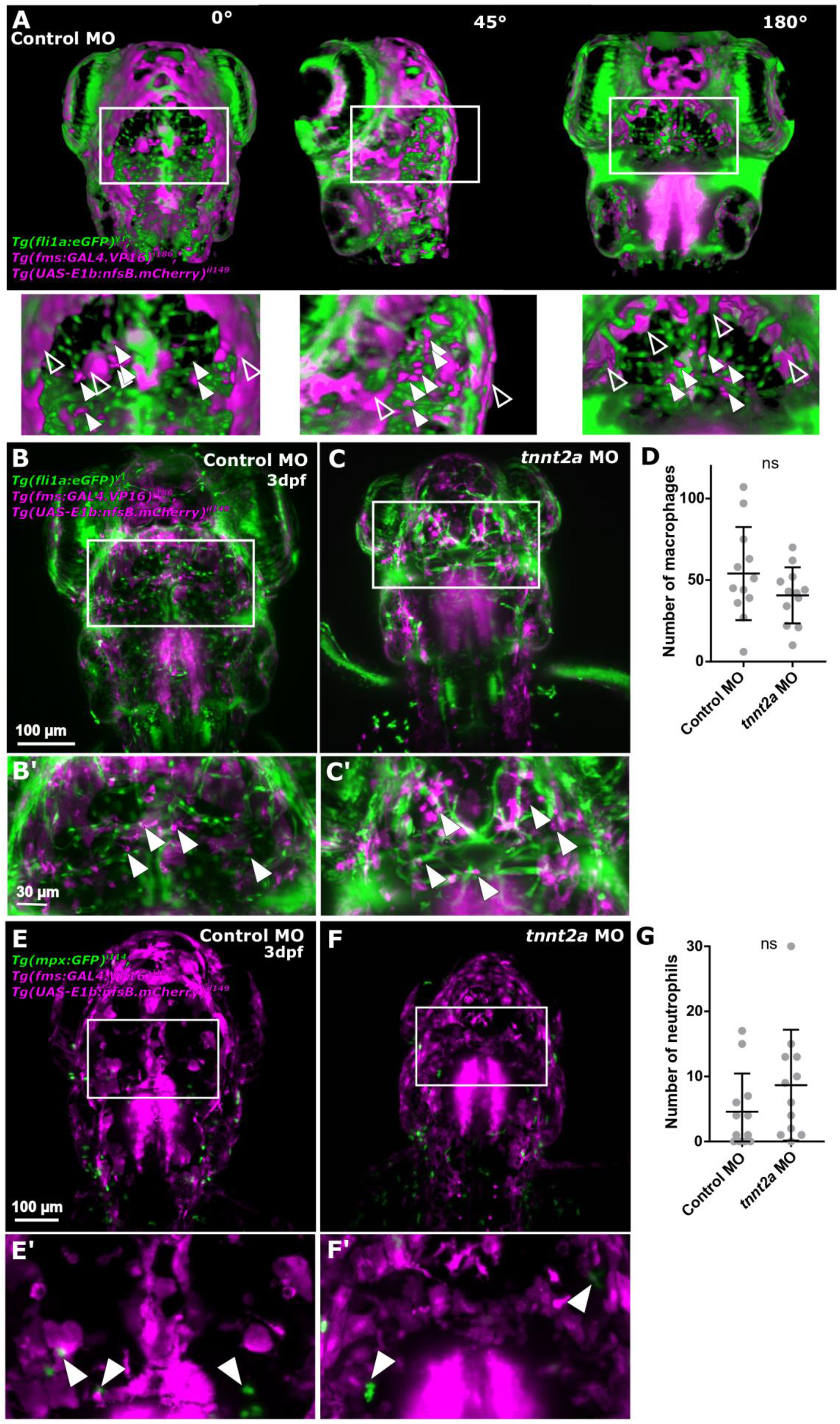
Lack of blood flow does not impact the number of cerebral innate immune cells. **(A)** Identification of immune cells was performed in 3D, allowing to discern non-specific signal (unfilled arrowhead) from immune cells (filled arrowhead). **(B-C)** Macrophages (magenta) were quantified examining the transgenic *Tg(fli1a:eGFP)*^*y1*^, *Tg(fms:GAL4.VP16)*^*i186*^, *Tg(UAS-E1b:nfsb.mCherry)*^*il149*^. **(D)** Number of macrophages was not changed upon blood flow loss (p 0.2356; n=12; 3dpf; Mann-Whitney U test). **(E-F)** Neutrophils (green) were examined in the transgenic reporter line *Tg(mpx:GFP)*^*i114*^, *Tg(fms:GAL4.VP16)*^*i186*^, *Tg(UAS-E1b:nfsb.mCherry)*^*il149*^. **(G)** Number of neutrophils was not changed upon blood flow loss (p 0.1708; n=12; 2 experimental repeats; 3dpf; Mann-Whitney U test).

To further examine whether the observed cell death was associated with altered tissue inflammation in the absence of altered macrophage and neutrophil numbers, we visualised the inflammatory mediator nitric oxide (NO) using the live dye DAF-FM. Visual assessment showed no difference in DAF-FM levels, thus inflammation, in the head (**Fig. 8A-C**) and trunk (**Fig. 8D-F**). This was confirmed by SNR quantification in the head (**Fig. 8G**; p 0.7967) and trunk (**Fig. 8H**; p 0.9371) which showed no differences comparing samples with blood flow to samples without blood flow.

**Figure 8.**
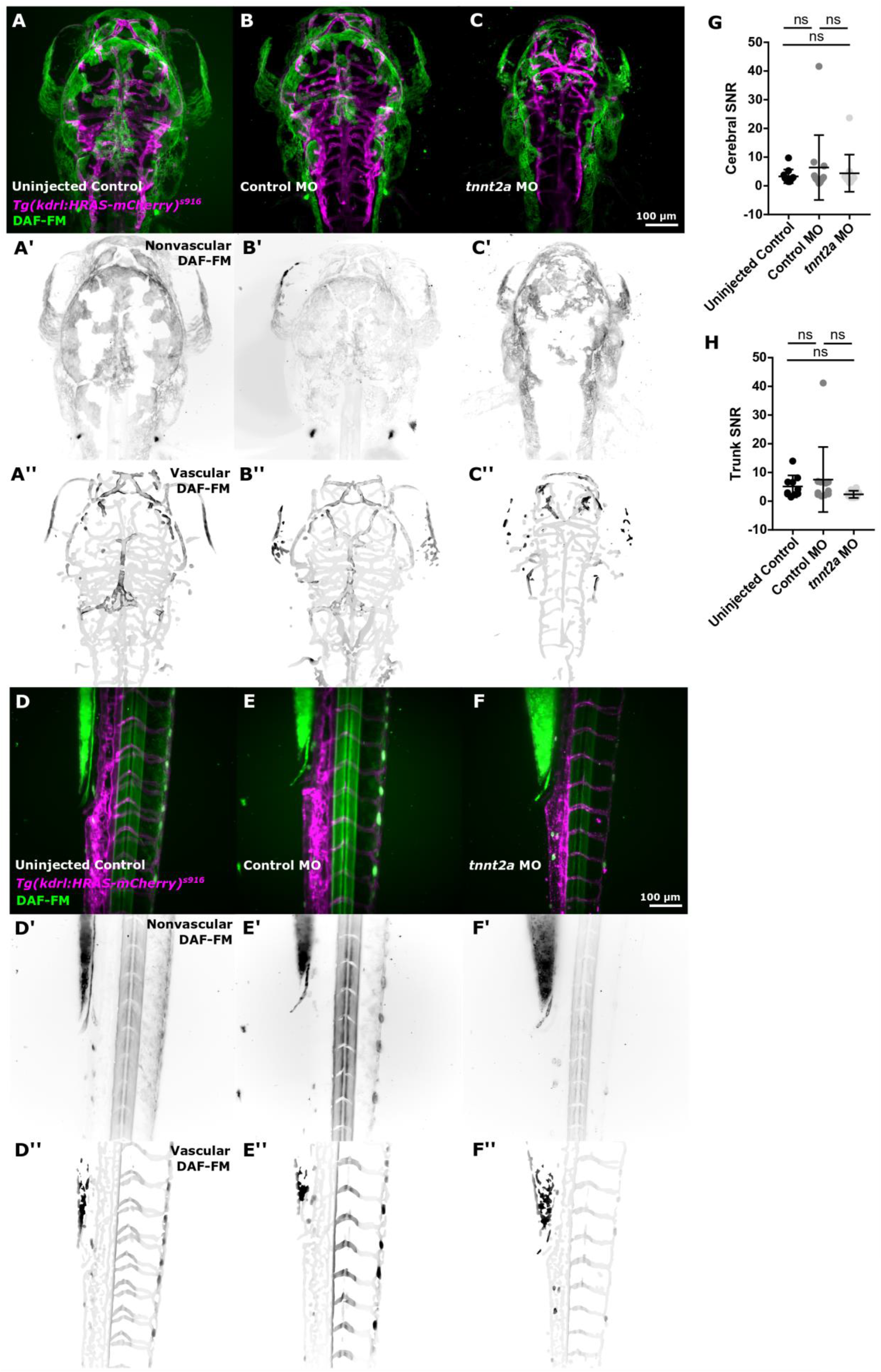
Inflammation is not increased by absent blood flow. **(A-C)** Examining nitric oxide (NO; visualized by DAF-FM) as an inflammatory marker, showed similar levels of NO in the head of uninjected controls, control MO, and *tnnt2a* MO (uninjected control n=11, control MO n=12, *tnnt2a* MO n=11; 3 experimental repeats; 3dpf). **(D-F)** DAF-FM levels in the trunk vasculature appeared visually similar between groups. **(G)** Quantification of cerebral DAF-FM SNR showed no difference between *tnnt2a* MO and uninjected controls (p>0.9999) or control MO (p>0.9999; Kruskal-Wallis test). **(H)** Quantification of trunk DAF-FM SNR showed no difference between *tnnt2a* MO and uninjected controls (p 0.1391) or control MO (p 0.1216; Kruskal-Wallis test).

Interestingly, we observed that DAF-FM signal, previously described to be localized in the bulbus arteriosus after 2dpf (Grimes et al., 2006), was absent in fish without blood flow (**Fig. S2**), suggesting bulbus arteriosus NO expression is blood flow dependent. Together, our data suggested that the lack of blood flow does increase cell death but not tissue inflammation at the investigated time-point.

## Discussion

In this study, we present the first assessment of the impact of blood flow on zebrafish embryonic vascular development from 2-to-5dpf and present the first comparison of the impact of flow in two vascular beds namely the head and trunk vasculature. We show that even though the overall response to lack of blood flow is similar in both vascular beds, the head vasculature is more severely impacted than trunk vasculature, with intra-cerebral vessels particularly being affected. Additionally, the effect of absent blood flow loss increases over time. Our data show that the lack of blood flow significantly increases cell death, and that this occurs without evidence of significant increase of tissue inflammation, as quantified by cerebral immune cell numbers and nitric oxide.

Our data showed that even in the complete absence of flow, sprouting angiogenesis can occur and stereotypic vascular patterning is preserved in the trunk and peripheral cerebral vessels. However, remodelling, such as intussusception in the CV is impaired as previously suggested (Karthik et al., 2018). The finding that central cerebral vessels show severely altered growth patterns, while peripheral vessels are less impacted, suggests that the response to blood flow is different in different vascular territories; whether this is due to a difference in vessel *formation* (e.g. peripheral vessels are formed from angiogenesis derived clusters, while central vessels are formed by angiogenesis from primary vessels) (Proulx et al., 2010; Siekmann et al., 2009) or *identity* (e.g. perineural, extra-cerebral, vs. intra-cerebral) (Vanhollebeke et al., 2015) requires future investigation.

Our work complements previous work (Serbanovic-Canic et al., 2017) which found blood flow cessation to induce EC apoptosis in zebrafish embryos at 30hpf, while our assessment of cerebral microglia numbers extends the examinations of Xu *et al*. who limited their studies to microglia of the tectum (Xu et al., 2016). Our finding that NO is lacking in the bulbus arteriosus at 3dpf in fish without blood flow is the first functional evidence of bulbus arteriosus NO to be blood flow dependent (Grimes et al., 2006).

Future studies might examine the nature of the detected apoptotic foci in the brain, and why these are predominantly found in the midbrain region. Our work focused on the contribution of cell death and our data suggest that apoptosis rather than necrosis to be the underlying mechanism. However, future work could entangle the contribution of apoptosis, necrosis, as well as proliferation.

We here presented the first comparisons, but the mechanisms of the observed effects in lacking blood flow, such as increased severity over time and different responses in vascular beds, require future investigation. This could include examining the expression of different mechanoreceptors and EC properties to complement previous findings about the importance and impact of blood flow (Feng Shuang et al., 2017; Mahmoud et al., 2017; Novodvorsky and Chico, 2014; Serbanovic-Canic et al., 2019; Souilhol et al., 2020; Watson et al., 2013). Similarly, molecular pathways and their context-dependent interpretation (such as VEGF, BMPs, Wnt (Benz et al., 2019; Liang et al., 2001; Vanhollebeke et al., 2015; Wiley and Jin, 2011)) might play a role in encountered differences in the head and trunk vasculature.

Together, our findings emphasize the differential response of ECs in different vascular beds to mechanical stimuli.

## Acknowledgments

We are grateful to Heba Ismail for feedback, Deepak Ailani for sharing chemicals, and Fiona Wright for technical support, as well as the Bateson Centre Zebrafish Facility Staff for support and advice. The authors thank all funders, including the University of Sheffield Department of Infection, Immunity and Cardiovascular Disease, Insigneo Institute for *in silico* Medicine, Medical Research Council, NC3Rs, and the British Heart Foundation.

## Declarations

### Author Contribution

Funding obtained by EK, PA, TC, JS-C, and PCE; Data Acquisition, EK, RS, GB, and KP; Investigation, Validation, and Data Curation, EK; Formal Visualization and Analysis, EK; Resources, PA, TC, JS-C, and PCE; Project Administration, EK and PA; Writing – Original Draft, EK; Writing – Review and Editing, all authors.

### Availability of data and material

Data are available upon request.

### Code availability

Code is available upon request.

### Ethics approval

Animal experiments were performed according to the rules and guidelines of institutional and UK Home Office regulations under the Home Office Project Licence 70/8588 held by TC.

### Consent for publication

Not applicable.

## Material and Methods

### Zebrafish Husbandry

Experiments were performed according to the rules and guidelines of institutional and UK Home Office regulations under the Home Office Project Licence 70/8588 held by TC.

Maintenance of adult zebrafish was performed as described in standard husbandry protocols (Aleström et al., 2019; Westerfield, 1993). Embryos, obtained from controlled mating, were kept in E3 (5mM NaCl, 0.17mM KCl, 0.33mM CaCl_2_, 0.33mM MgSO_4_) medium buffer with methylene blue and staged according to Kimmel *et al*. (Kimmel et al., 1995). The following transgenic reporter lines were used: *Tg(kdrl:HRAS-mCherry)*^*s916*^ (Chi et al., 2008) visualizes EC membrane, *Tg(fli1a:eGFP)*^*y1*^ (Lawson and Weinstein, 2002) visualizes EC cytosol, *Tg(flk1:nls-eGFP)*^*zf109*^ (Blum et al., 2008) visualizes EC nuclei, *Tg(mpx:GFP)*^*i114*^ (Renshaw et al., 2006) visualizes neutrophils, and *Tg(fms:GAL4.VP16)*^*i186*^, *Tg(UAS-E1b:nfsB.mCherry)*^*il149*^ (Gray et al., 2011) visualizes macrophages.

### Morpholino injection

Development of functional heart contraction was inhibited via injection of *tnnt2a* ATG morpholino (1.56 ng final concentration), as described in (Sehnert and Stainier, 2002; Sehnert et al., 2002) (sequence 5’-CATGTTTGCTCT GATCTGACACGCA-3’). Control morpholino injections (5’-CCTCTTACCTCAGTTATTTATA-3’; no target sequence and little/no biological activity; Genetools, LLC) were performed with the above final concentration to study off-target effects of injections. Injections were conducted at one-cell-stage using phenol red as injection tracer.

### Chemical and histological stains

*In vivo* visualization of cell death was performed using 2 μg/mL solution of Acridine Orange (Sigma) in 1X E3 for 2h in 3dpf embryos, followed by 3 washes in E3 before image acquisition (Verduzco and Amatruda, 2011). *In vivo* visualization of inflammation via nitric oxide (NO) (Zhang et al., 2019) was performed using 2.5μM DAF-FM-DA (Molecular Probes; D23844) (Kojima et al., 1998) for 6h in 3dpf embryos. DMSO control was performed at the same concentration and duration.

### Image Acquisition Settings and Properties

Anaesthetized embryos were embedded in 2% LMP-agarose with 0.01% Tricaine in E3 (MS-222, Sigma). Data acquisition of the cranial and trunk vasculature was performed using a Zeiss Z.1 light sheet microscope, Plan-Apochromat 20×/1.0 Corr nd=1.38 objective, dual-side illumination with online fusion, activated Pivot Scan, image acquisition chamber incubation at 28°C, and a scientific complementary metal-oxide semiconductor (sCMOS) detection unit. The properties of acquired data were as follows: 16bit image depth, 1920 × 1920 × 400-600 voxel (x,y,z; approximate voxel size of 0.33 × 0.33 × 0.5 μm, respectively). Multicolour images in double-transgenic embryos were acquired in sequential mode.

### Image Analysis

As *tnnt2a* can develop considerable oedema, we ensured comparability between samples by standardized image acquisition from the dorsal view, including the most dorsal vessel the dorsal longitudinal vein (DLV) to the more ventral basilar artery (BA; **Fig. S1A**). For image analysis ROI selection was performed as previously described (**Fig. S1B**) (Kugler et al., 2019a).

#### Nuclei detection

6-by-6 neighbourhood median filter (Lim, 1990) to remove salt-and-pepper noise (Gonzalez and Woods, 2007) and background removal with the rolling ball algorithm with size 200 (Sternberg, 1983), 2D maximum intensity projection, and detection of local noise maxima using Fiji Software (Burger and Burge, 2008; Schindelin et al., 2012). Voronoi diagram was established following Otsu thresholding (Otsu, 1979).

##### Signal intensities

Signal intensity measurement of Acridine Orange and DAF-FM were conducted by creating 3D vascular masks following Sato filter for vascular enhancement and Otsu thresholding segmentation as previously described (Kugler et al., 2018, 2019a, 2020). Signal mean was quantified in ROIs in 2D MIPs. Signal-to-noise (SNR) ratio was quantified as mean signal in ROI divided by the standard deviation of the background (ROI placed outside the fish with a size of 10μm x10μm). Acridine orange foci were detected in ROI following 2D Median filtering using detection of maxima with an intensity over 10.

#### Manual analysis

All analysis was performed using Fiji (Schindelin et al., 2012). Diameter of basilar artery (BA) was measured approximately 30μm before bifurcating into posterior (caudal) communicating segments (PCS).

Posterior cerebral vein (PCeV) was measured approximately 20μm before turning dorsally.

Posterior cardinal vein (PCV) and dorsal aorta (DA) diameters were measured above the cloaca, with three measurement points each in the same animal.

Intersegmental vessel (ISV) lengths of arterial (aISVs) and venous (vISVs) were measured above the cloaca, with three measurement points of the same vessels each in the same animal.

#### Immune cells

Intracranial macrophages and neutrophils were quantified manually in 3D after ROI selection of the dorsal cerebral vasculature.

### Statistics and Data Representation

Gaussian distribution conformation was evaluated using the D’Agostino-Pearson omnibus test [21]. Statistical analysis was performed using One-way ANOVA or paired Students t-test in GraphPad Prism Version 7 (GraphPad Software, La Jolla California USA). Statistical significance was represented as: p<0.05 *, p<0.01 **, p<0.001 ***, p<0.0001 ****. Graphs show mean values ± standard deviation unless otherwise indicated. Image representation and visualization was done with Inkscape Version 0.48 (https://www.inkscape.org). Images were visualized as maximum intensity projections (MIPs). 3D rendering was performed using Arivis software (arivis AG, Munich, Germany).

## Supplementary Material

**Fig S1:**
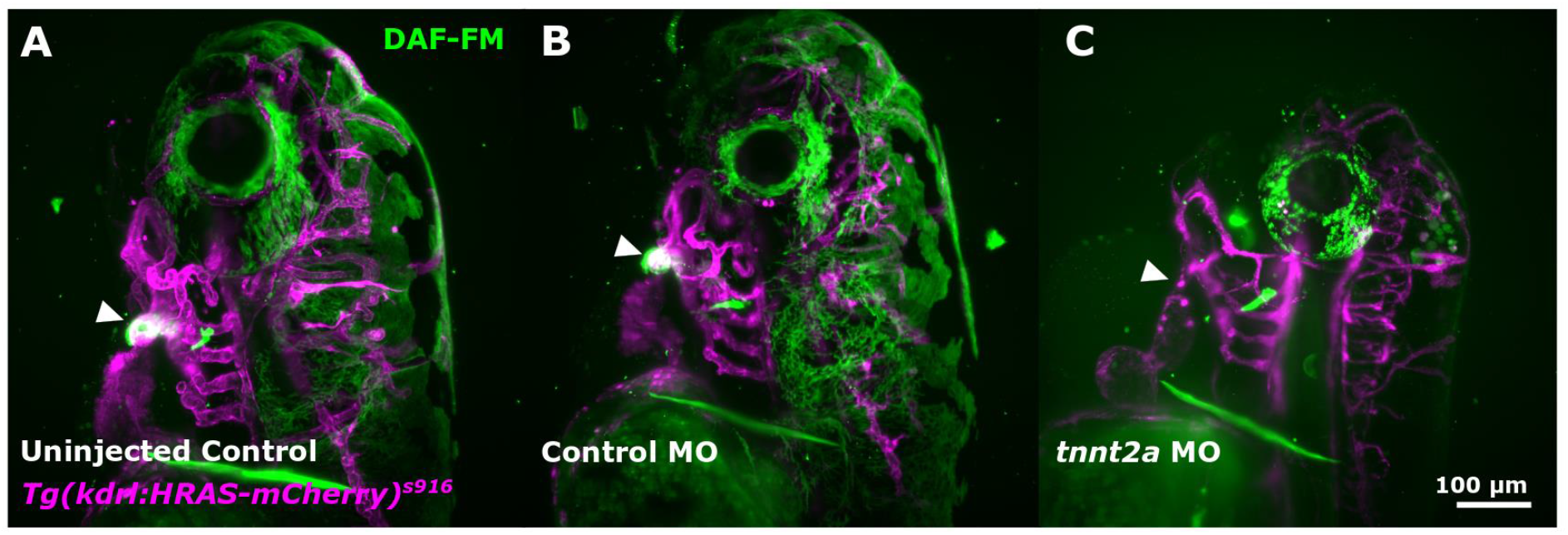
Cardiac NO is absent in animals without cardiac contraction. DAF-FM signal in bulbus arteriosus (white arrowheads) was lacking in *tnnt2a* MO (uninjected control n=11, control MO n=12, *tnnt2a* MO n=11; 3 experimental repeats; 3dpf).

**Figure S2:**
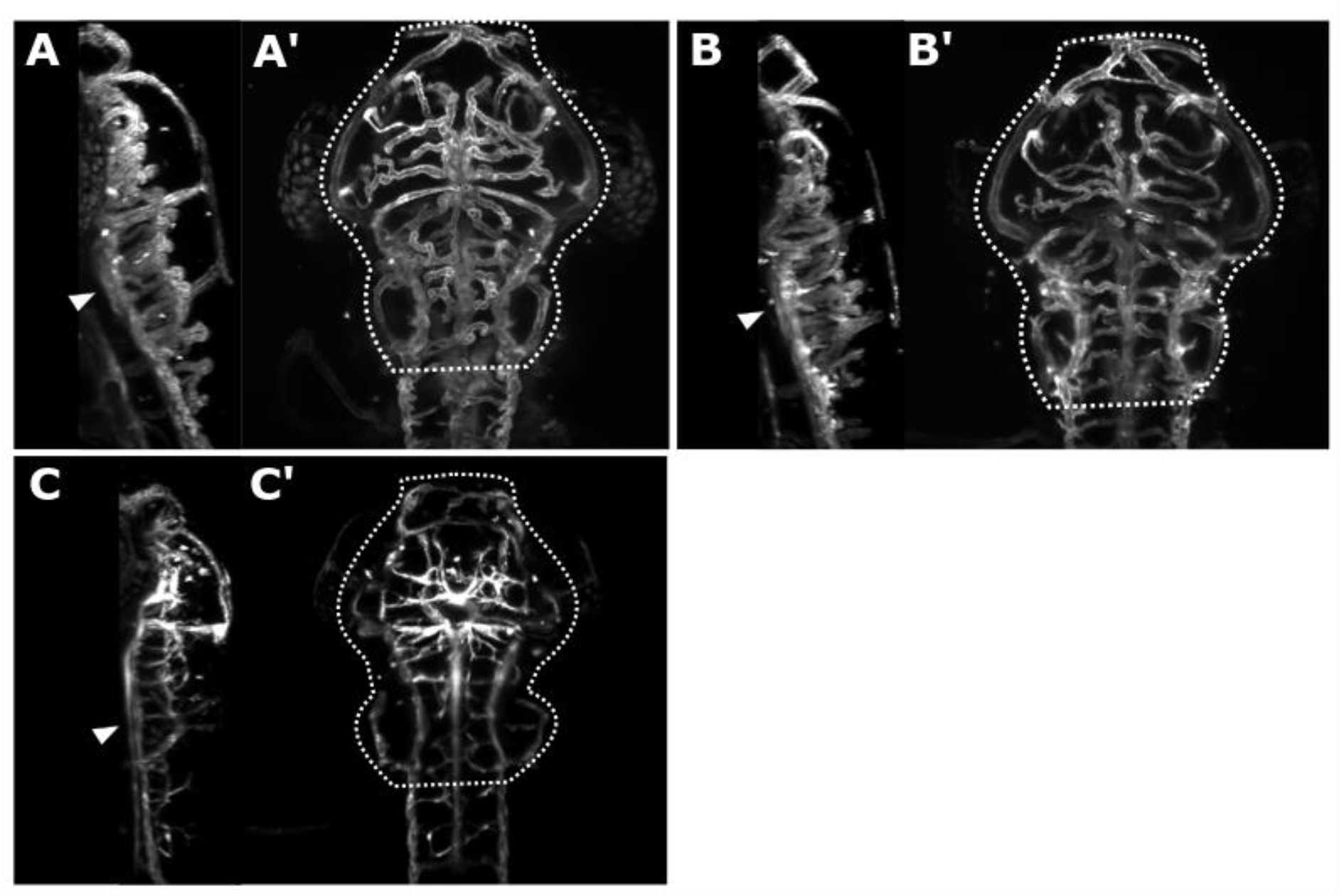
ROI to compare samples.

